# Drivers of Vessel Progenitor Fate Define Intermediate Mesoderm Dimensions by Inhibiting Kidney Progenitor Specification

**DOI:** 10.1101/2024.02.22.581649

**Authors:** Elliot A. Perens, Deborah Yelon

## Abstract

Proper organ formation depends on the precise delineation of organ territories containing defined numbers of progenitor cells. Kidney progenitors reside in bilateral stripes of posterior mesoderm that are referred to as the intermediate mesoderm (IM). Previously, we showed that the transcription factors Hand2 and Osr1 act to strike a balance between the specification of the kidney progenitors in the IM and the vessel progenitors in the laterally adjacent territory. Recently, the transcription factor Npas4l – an early and essential driver of vessel and blood progenitor formation – was shown to inhibit kidney development. Here we demonstrate how kidney progenitor specification is coordinated by *hand2*, *osr1*, and *npas4l*. We find that *npas4l* is necessary to inhibit IM formation. Consistent with the expression of *npas4l* flanking the medial and lateral sides of the IM, our findings suggest roles for *npas4l* in defining the IM boundaries at each of these borders. At the lateral IM border, *hand2* promotes and *osr1* inhibits the formation of *npas4l*-expressing lateral vessel progenitors, and *hand2* requires *npas4l* to inhibit IM formation and to promote vessel formation. Meanwhile, *npas4l* appears to have an additional role in suppressing IM fate at the medial border: *npas4l* loss-of-function enhances *hand2* mutant IM defects and results in excess IM generated outside of the lateral *hand2*-expressing territory. Together, our findings reveal that establishment of the medial and lateral boundaries of the IM requires inhibition of kidney progenitor specification by the neighboring drivers of vessel progenitor fate.

## INTRODUCTION

Organ development commences with the establishment of well-defined organ territories characterized by unique gene expression signatures. Neighboring organ territories often abut one another, and the proximity of adjacent territories necessitates coordination over time to refine gene expression and ensure precise boundaries between domains (Briscoe & Small, 2015). This specification process can depend on reciprocal interactions in which the promotion of the proper cell fate coincides with the repression of alternative fates (Davidson, 2010).

The vertebrate posterior mesoderm is composed of neighboring organ territories arranged along the medial-lateral axis (Prummel et al., 2020). One such territory positioned within the posterior mesoderm is the intermediate mesoderm (IM), a pair of bilateral stripes of kidney and urinary tract progenitors (Dressler, 2009; Saxen, 1987). The IM is often defined by the expression of transcription factor genes such as *pax2*, *lhx1*/*lim1*, and *wt1*, which are expressed in and required for IM and kidney development (Bollig et al., 2006; Dressler & Douglass, 1992; Krauss et al., 1991; Kreidberg et al., 1993; Majumdar et al., 2000; Serluca & Fishman, 2001; Toyama & Dawid, 1997). Induction of IM formation is dependent on neighboring tissues, including surface ectoderm and paraxial mesoderm, and morphogens, including BMP, FGF, and Wnt, although the precise connections between the morphogens and their sources are not well established (James & Schultheiss, 2003, 2005; Mauch et al., 2000; Naylor et al., 2017; Obara-Ishihara et al., 1999; Tsang et al., 2000; Warga et al., 2013). Furthermore, it remains unclear how the dimensions of the IM are determined. In particular, the mechanisms that establish the well delineated gene expression boundary between the IM and the adjacent vessel progenitor populations have not been determined. Some prior studies have hinted at antagonistic developmental relationships between transcription factors that establish the IM and vessel lineages. For example, overexpression of the vascular and hematopoietic transcription factor genes *tal1* and *lmo2* induces ectopic vessel and blood formation while inhibiting IM specification (Gering et al., 2003). Conversely, loss of *pax8* function can result in ectopic vessel progenitor marker expression in the place of kidney progenitor formation (Buisson et al., 2015). These findings suggest roles for these transcription factors as repressors of alternative fates, ensuring precise gene expression boundaries between the neighboring IM and vessel progenitor territories.

Previously, we and others identified roles for the transcription factors Hand2 and Osr1 in coordinating kidney and vessel progenitor fates in the zebrafish embryo (Drummond et al., 2022; Mudumana et al., 2008; Perens et al., 2021; Perens et al., 2016). In the zebrafish posterior mesoderm, the IM is initially flanked by two territories: a lateral territory that expresses *hand2* and *osr1*, and a medial territory that contains vessel and blood progenitors (Mudumana et al., 2008; Perens et al., 2016). Later, a separate set of lateral vessel progenitors (LVPs) arises between the IM and the *hand2*- and *osr1*-expressing territory (Perens et al., 2016). *hand2* is required to limit the lateral extent of the IM by inhibiting ectopic IM gene expression within the *hand2*-expressing territory, and *hand2* is also required to promote LVP fate (Perens et al., 2016). Conversely, *osr1* is required for generating the full complement of IM while also inhibiting the premature emergence of LVPs (Perens et al., 2021). *hand2*;*osr1* double mutants form IM and LVPs at levels comparable to wild-type, indicating an antagonistic genetic relationship between these two genes and potential coupling of these two cell fates (Perens et al., 2021). Together, these findings led to a model in which *hand2* and *osr1* work in opposition to each other at the lateral border of the developing IM to ensure the proper allocation of IM and LVP cell fates in this territory.

More recently, Mattonet and colleagues found additional factors that promote vessel progenitor formation and also inhibit kidney cell fate (Mattonet et al., 2022). Specifically, they found that *npas4l* (*cloche*), an early essential regulator of vessel and blood progenitor formation (Liao et al., 1998; Liao et al., 1997; Reischauer et al., 2016; Stainier et al., 1995; Sumanas et al., 2005), along with downstream vessel-promoting factors *tal1* and *lmo2*, inhibits kidney formation. Notably, they determined that *npas4l*-expressing cells, when lacking *npas4l* function, contribute to the pronephron (the zebrafish embryonic kidney), instead of the vascular and hematopoietic systems. How *hand2*, *osr1*, and *npas4l* function together to coordinate vessel progenitor and kidney fates, however, is unknown. In particular, does *npas4l* simply act downstream of *hand2* and *osr1* to regulate LVP and pronephron progenitor fates emerging from the lateral IM border, or is there a broader requirement for vessel-promoting factors to repress kidney fate in the posterior mesoderm?

Here, through genetic analyses and molecular anatomical studies, we assess how IM and vessel progenitor fates are coordinated by *hand2*, *osr1*, and *npas4l*. First, we find that *hand2* and *osr1* regulate the formation of *npas4l*-expressing LVPs and that *hand2* requires *npas4l* both to inhibit kidney and to promote vessel progenitor formation, consistent with *npas4l* functioning downstream of *hand2* and *osr1* to regulate LVP and kidney fates at the lateral IM border. On the other hand, we find that, unlike *hand2*, *npas4l* is necessary to inhibit IM formation outside of the *hand2*-expressing territory, and loss of *npas4l* function enhances IM defects in *hand2* mutants, suggesting an additional role for *npas4l* distinct from *hand2*. Taken together, these data support a model in which *npas4l* inhibits IM fate at both the medial and lateral borders of the IM. More broadly, our findings highlight that the dimensions of the IM and the boundaries between adjacent vessel and kidney organ territories are established by the inhibition of kidney fate by neighboring vessel-promoting factors.

## MATERIALS AND METHODS

### Zebrafish

We generated embryos by breeding wild-type zebrafish, heterozygotes for the *osr1* mutant allele *osr1^el593^* (RRID: ZFIN_ZDB-ALT-171010-14) (Askary et al., 2017), heterozygotes for the *hand2* mutant allele *han^s6^* (RRID: ZFIN_ZDB-GENO-071003-2) (Yelon et al., 2000), heterozygotes for the *npas4l* mutant allele *npas4l^s5^*(RRID: ZDB-FISH-150901-21185) (Field et al., 2003), carriers of the transgene *Tg*(*hand2:egfp*)*^pd24^* (RRID: ZFIN_ZDB-GENO-110128-35) (Kikuchi et al., 2011), carriers of the transgene *Tg*(*hsp70:osr1-t2A-BFP*)*^sd63^*(RRID: ZFIN_ZDB-TGCONSTRCT-221201-1) (Perens et al., 2021), and carriers of the transgene *Tg(hsp70:FLAG-hand2-2A-mCherry*)*^sd28^* (RRID: ZFIN_ZDB-GENO-141031-2) (Schindler et al., 2014). All zebrafish work followed protocols approved by the UCSD IACUC.

### Heat Shock

For induction of heat shock-regulated expression, embryos were exposed to 37°C for one hour and then returned to 28°C. Heat shocks were performed at tailbud stage. Following heat shock, transgenic embryos were distinguished by their BFP or mCherry fluorescence. Nontransgenic embryos served as controls.

### Genotyping

PCR genotyping was conducted as previously described for *osr1^el593^*mutants (Askary et al., 2017), *han^s6^*mutants (Yelon et al., 2000), and *han^s6^* mutants containing *Tg(hand2:EGFP)^pd24^* (Perens et al., 2016). *npas4l^s5^*genotyping was conducted using the forward primer 5′-TCTGCCGGGGTTTATCGTT-3′ and reverse primer 5′-ACTCGTGTACGTTCTCAGACAG-3′. The reverse primer creates a mismatch that generates a HpyCH4III restriction site in the *npas4l^s5^* allele amplicon, while the wild-type allele amplicon is uncut.

### Injection

We synthesized capped mRNA from a pCS2+-*npas4l* plasmid (Reischauer et al., 2016) using the Ambion mMESSAGE mMACHINE kit, and we injected 5-14.5 ng mRNA into embryos at the one-cell stage.

### *In situ* hybridization

Standard whole-mount *in situ* hybridization were performed as previously described (Thomas et al., 2008) using the following probes: *atp1a1a.4* (ZDB-GENE-001212-4), *cdh17* (ZDB-GENE-030910-3), *etv2* (*etsrp*; ZDB-GENE-050622-14), *hand2* (ZDB-GENE-000511-1), *lhx1a* (*lim1*; ZDB-GENE-980526-347), *npas4l* (*clo*; ZDB-GENE-070117-2515), *osr1* (ZDB-GENE-070321-1), *pax2a* (ZDB-GENE-990415-8), *wt1b* (ZDB-GENE-050420-319). RNA antisense probes were generated using cDNA templates for all probes except *npas4l*, which was generated using a PCR template (Reischauer et al., 2016).

### Immunofluorescence and Hybridization Chain Reaction

Whole-mount immunofluorescence was performed as previously described (Cooke et al., 2005), using polyclonal antibodies against Pax2a at 1:100 dilution (Genetex, GTX128127) (RRID: AB_2630322) and against GFP at 1:100 dilution (Life Technologies, A10262) (RRID: AB_2534023), and the secondary antibodies goat anti-chick Alexa Fluor 488 (Life Technologies, A11039) (RRID: AB_2534096) and goat anti-rabbit Alexa Fluor 594 (Life Technologies, A11012) (RRID: AB_10562717), both at 1:100 dilution. Samples were then placed in SlowFade Gold anti-fade reagent (Life Technologies) and mounted in 50% glycerol in PBST.

Whole mount hybridization chain reaction (HCR) fluorescent *in situ* hybridization (FISH) was performed based on Multiplexed HCR v3.0 protocol from Molecular Instruments, Inc. as described in (Choi et al., 2018), followed by immunohistochemistry. For HCR FISH, probes and amplifiers were obtained from Molecular Instruments, Inc. Probes were generated based on sequences for *npas4l* (NCBI Reference Sequence: NM_001329912.1, Lot # RTI256) and *hand2* (NCBI Reference Sequence: NM_131626.3, Lot # RTB193). Probe-amplifier combinations used were *npas4l* with B3 Alexa 488 and *hand2* with B1 Alexa 647. Briefly, embryos were fixed in 4% paraformaldehyde overnight, dehydrated in methanol overnight, rehydrated into PBS + 0.1% Tween 20 (PBST), permeabilized with 100% acetone for 12 minutes, washed in PBST, pre-hybridized with Hybridization Buffer (Molecular Instruments, Inc.) for one hour at 37°C, and incubated with probes overnight at 37°C using 10 pmol probe in 100ul Hybridization Buffer overnight. Embryos were washed in Wash Buffer (Molecular Instruments, Inc.) and 5X SSC + 0.1% Tween. 20 (SSCT). For amplification, embryos were pre-incubated in Amplification Buffer (Molecular Instruments, Inc.) for one hour and then incubated with heat activated hairpins (5 ul hairpin per 250 ul Amplification Buffer) all at room temperature. The following day, embryos were washed in 5X SSCT followed by PBST washes. Then, for immunohistochemistry, embryos were placed in block (PBST + 0.1% Triton X100 (PBSTT) + 2% sheep serum) for two hours at room temperature before incubating with Pax2a antibody, as above, overnight at 4°C. Embryos were washed several times with PBSTT and re-blocked in PBSTT with 1% sheep serum before incubating with secondary antibody, as above, overnight at 4°C. Last, embryos were washed in PBSTT, briefly incubated in SlowFade,Gold reagent and stored and mounted in 50% glycerol in PBST.

### Imaging

Bright-field images were captured with a Zeiss Axiocam on a Zeiss Axiozoom microscope and processed using Zeiss AxioVision. Confocal images were collected by Leica SP5 (Figs. 5 and 6) and SP8 (Fig. 1) confocal laser-scanning microscopes using a 10× dry objective. Slice thicknesses were 1 µm (Figs. 5 and 6) and “System Optimized” based on Leica LAS X software (Fig. 1). Images were analyzed using Imaris software (Bitplane).

**Figure 1.**
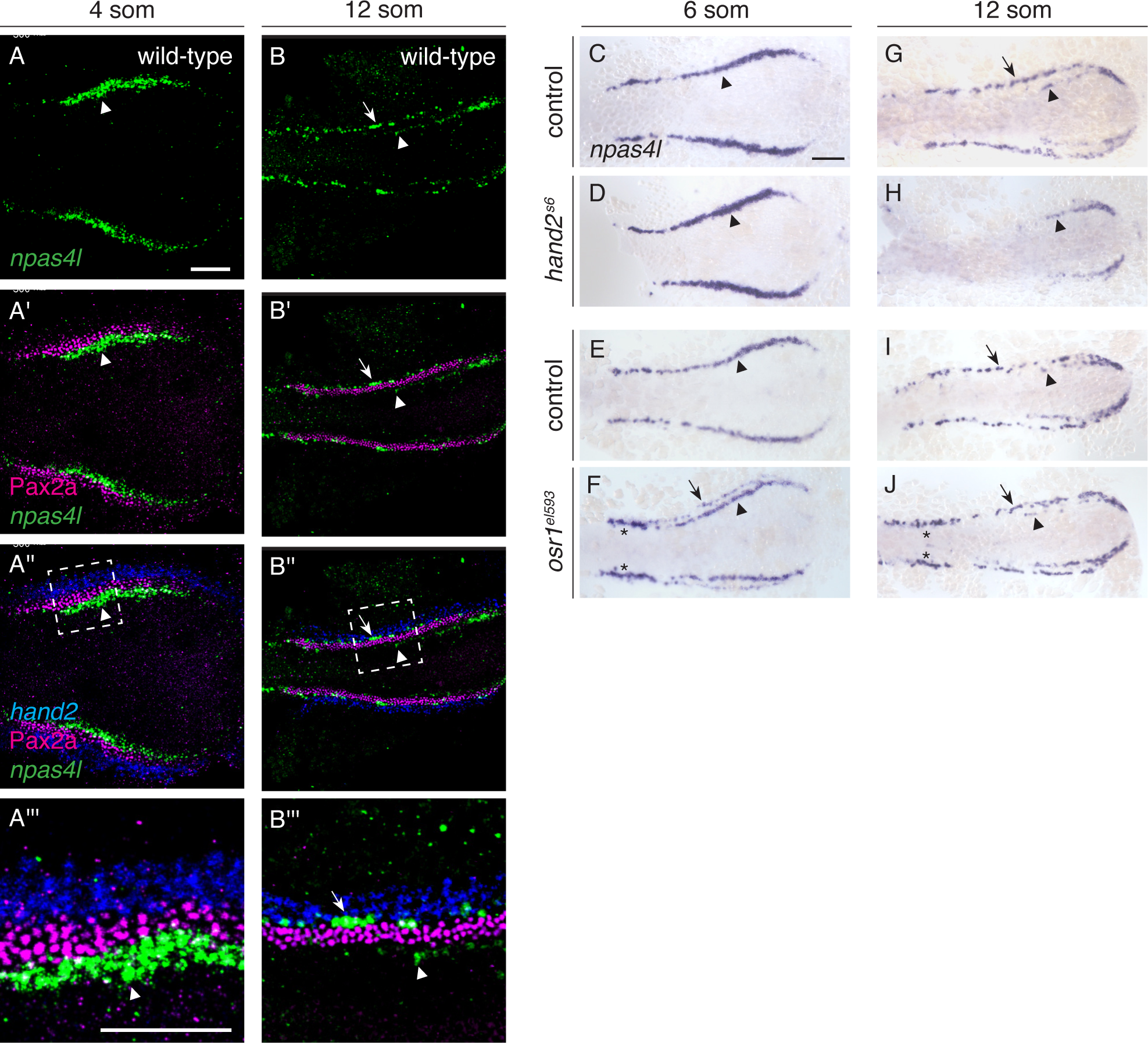
*hand2* and *osr1* regulate the development of *npas4l*-expressing lateral vessel progenitors. (A,B) *In situ* hybridization chain reaction (*npas4l*, *hand2*) and immunohistochemistry (Pax2a) label components of the wild-type posterior mesoderm in dorsal views, anterior to the left, of three-dimensional reconstructions. (A’’’, B’’’) Magnification of boxed regions in A’ and B’. (A-A’’’) At the 4 somite stage (som), *npas4l* is expressed in bilateral territories (green, arrowheads) medial to the IM marker Pax2a (magenta) and *hand2* (blue). (B-B’’’) At 12 som, *npas4l*-expressing lateral vessel progenitors (arrows) have arisen lateral to Pax2a and medial to *hand2*. Faint *npas4l* expression remains in the medial territory (arrowheads). (C-J) *In situ* hybridization shows *npas4l* expression in wild-type control (C,G,E,I), *hand2^s6^* mutant (D,H) and *osr1^el593^* mutant (F,J) embryos; dorsal views, anterior to the left, at 6 som (C-F) and 12 som (G-J). (C-F) At 6 som, *npas4l* is expressed in a relatively medial territory (arrowhead) on each side of the wild-type (C,E), *hand2^s6^* (D), and *osr1^el593^* (F) posterior mesoderm. In *osr1^el593^*mutants, *npas4l* is also expressed in relatively lateral cells (F, arrow), and its expression is increased in a distinct proximal region (F, asterisks). Lateral *npas4l* expression was observed in 84% of *osr1^el593^* mutants (n=31) and 2% of wild-type control siblings (n=110); furthermore, the lateral expression observed in wild-type embryos was restricted to only a few cells at the anterior extent of the posterior mesoderm (Fig. S1C). Similarly premature expression of *etv2* in lateral cells was seen in *osr1* mutants previously (Perens et al., 2021). (G-J) At 12 som, *npas4l* is expressed in a relatively lateral territory (arrows) on each side of wild-type (G,I) and *osr1^el593^* mutant (J) embryos; in *osr1^el593^* embryos, *npas4l* expression remains increased proximally (J, asterisks). In *hand2^s6^* mutants (H), *npas4l* is not expressed in this lateral territory (100%, n=20). At this stage, a small amount of *npas4l* expression is still observed in the posterior medial vessel progenitors in *hand2^s6^*mutants (H, arrowhead). n=77 (C), 30 (D), 110 (E), 31 (F), 66 (G), 20 (H), 53 (I), 18 (J). Scale bars: 100 μm.

### Cell counting

Pax2a^+^ and GFP^+^ cells were counted as described previously (Perens et al., 2021). Briefly, we removed the yolk and the anterior region before flat-mounting and imaging the embryos. In each embryo, we analyzed three-dimensional reconstructions and individual optical sections of a contiguous, representative region measuring 250 µm in length. Pax2a^+^ cells were differentiated from background immunofluorescence by decreasing brightness in Imaris until staining clearly outside of IM was no longer visible.

### Statistics and replication

Statistical analysis was performed using Graphpad Prism 8 to conduct non-parametric Mann–Whitney *U*-tests. All results represent at least two independent experiments (technical replicates) in which multiple embryos, from multiple independent matings, were analyzed (biological replicates). n values in figure legends represent number of embryos with corresponding genotype that were analyzed. In cases in which there were notable variations in phenotype, specific information is detailed in the figure legends.

## RESULTS

### *npas4l* is expressed at the medial and lateral borders of the intermediate mesoderm

To gain insight into how progenitor fates in the posterior mesoderm are coordinated by *npas4l*, *hand2* and *osr1*, we first examined the spatial and temporal characteristics of *npas4l* expression relative to other landmarks in the posterior mesoderm. Previously, Reischauer and colleagues showed that *npas4l* is initially expressed in bilateral regions of the posterior mesoderm as early as the tailbud stage (Reischauer et al., 2016). We found this expression to be immediately medial to the IM (Fig. 1A). At approximately the 8 somite stage (som), a second, more lateral population of *npas4l*-expressing cells arises bilaterally along the lateral border of the IM, at the interface between the IM and the *hand2*-expressing territory (Fig. 1B; Fig. S1A-E). Thus, as found previously for *etv2* and *tal1* (Kohli et al., 2013; Perens et al., 2016), in the posterior mesoderm, *npas4l* is expressed bilaterally in two stripes of expression that flank the medial and lateral aspects of the IM.

The temporal dynamics of *npas4l* expression in the posterior mesoderm, however, differ from those of *etv2* and *tal1*. Previously, *npas4l* expression in the posterior mesoderm was shown to be expressed at tailbud, before *etv2* and *tal1*, which initiate expression at 2 som (Reischauer et al., 2016). After that, *npas4l* expression is transient in these medial vessel progenitors (Fig. S1A-E). *npas4l* expression begins to decline at the anterior aspect of the medial vessel progenitor expression territory starting at 8 som (Fig. S1D) and remains expressed only in the most posterior of the medial vessel progenitors by 12 som (Fig. 1B,G,I; Fig. S1E). In contrast, expression of *etv2* persists in the medial vessel progenitors until at least 13 som when these progenitors begin to migrate to the midline (Fig. S1F-J). Similarly, *npas4l* expression in the LVPs precedes *etv2* and *tal1* expression. More specifically, in wild-type embryos, *etv2* and *tal1* expression in LVPs was usually not observed until 11 som (Fig. S1F-J; (Perens et al., 2016)); in contrast, *npas4l* expression in the LVPs can sometimes be seen as early as 6 som in a small number of the most anterior LVPs (Fig. S1C) and is observed throughout most of the LVPs by 8 som (Fig. S1D). Finally, at 18 som, *npas4l* expression is globally decreased while *etv2* and *tal1* expression levels remain persistently high (Reischauer et al., 2016). These findings are consistent with *npas4l* functioning transiently to initiate vessel progenitor specification prior to the expression of *etv2* and *tal1*.

### *hand2* is required for formation of *npas4l*-expressing lateral vessel progenitors while *osr1* inhibits their premature emergence

Previously, we concluded that *hand2* and *osr1* control LVP formation based on our observations of *etv2* and *tal1* expression in the LVPs (Perens et al., 2021; Perens et al., 2016). For example, we found that *hand2* is required for expression of *etv2* and *tal1* in LVPs (Perens et al., 2016). Because *npas4l* expression precedes *etv2* and *tal1* expression, we wondered whether *hand2* has the same effect on *npas4l* expression in LVPs. As with *etv2* and *tal1* expression, we did not observe *npas4l* expression in LVPs in *hand2* mutants (Fig. 1H), consistent with LVPs failing to form in the absence of *hand2* function. Conversely, while *osr1* mutants ultimately form the correct quantity of LVPs, we have shown that LVP expression of *etv2* initiates prematurely in *osr1* mutants: in wild-type embryos, most LVPs express *etv2* by 11 som, whereas in *osr1* mutants, *etv2* expression was observed as early as 6 som in some LVPs and most LVPs expressed *etv2* by 8 som (Perens et al., 2021). Similarly, we observed the premature appearance of *npas4l*-expressing LVPs in *osr1* mutants. Specifically, while *npas4l* LVP expression was rarely observed prior to 8 som in wild-type embryos (Fig. S1A-E), some *osr1* mutants exhibited *npas4l* expression in LVPs by 4 som (n=4/17 *osr1* mutants; n=0/57 wild-type controls), and the majority of *osr1* mutants expressed *npas4l* throughout the LVPs by 6 som (Fig. 1F; n=7/10 *osr1* mutants; n=0/36 wild-type controls). In contrast, the expression of *npas4l* in medial vessel progenitors did not appear altered in *hand2* or *osr1* mutants (Fig. 1C-J).

Consistent with a role for *hand2* upstream of *npas4l* in LVPs, *hand2* overexpression increased the number of *npas4l*-expressing cells (Fig. S2A,B). This *hand2* overexpression phenotype was similar to that observed based on *etv2* and *tal1* expression (Perens et al., 2016). In contrast, *osr1* overexpression inhibited *npas4l* expression in LVPs (Fig. S2C,D), also similar to the pattern observed with *etv2* expression (Perens et al., 2021). Last, *hand2* and *osr1* expression appeared unaltered in *npas4l* mutants (Fig. S3). Altogether, these findings suggest that *hand2* and *osr1* act upstream of *npas4l* expression to regulate the formation of LVPs.

### Inhibition of pronephron formation by *hand2* requires *npas4l* function

Because *hand2* is required for formation of *npas4l*-expressing LVPs, we next wanted to determine whether *hand2* regulation of IM and LVP formation is dependent on *npas4l*. Previously, we found that overexpression of *hand2* caused robust inhibition of pronephron development and promoted ectopic vessel progenitor formation (Perens et al., 2016). Interestingly, the effect of *hand2* overexpression on pronephron formation was greatly blunted in the absence of *npas4l* function. That is, although some of the pronephron often failed to form when *hand2* was overexpressed in *npas4l* mutants, the inhibitory effect of *hand2* overexpression was milder than in wild-type siblings. (Fig. 2A-D). Not surprisingly, based on the essential role of *npas4l* in vessel progenitor formation (Liao et al., 1998; Liao et al., 1997; Stainier et al., 1995; Sumanas et al., 2005), *hand2* overexpression could not promote vessel formation in the absence of *npas4l*, as indicated by the scant *etv2* expression in vessel progenitors and *flk1* expression in the vasculature (Fig. 2E-L). Together, these findings suggest that *npas4l* is required downstream of *hand2* both to inhibit pronephron formation and to promote vessel formation. Additionally, these findings are consistent with a model in which *hand2* inhibition of pronephron formation is dependent on its ability to promote vessel progenitor formation.

**Figure 2.**
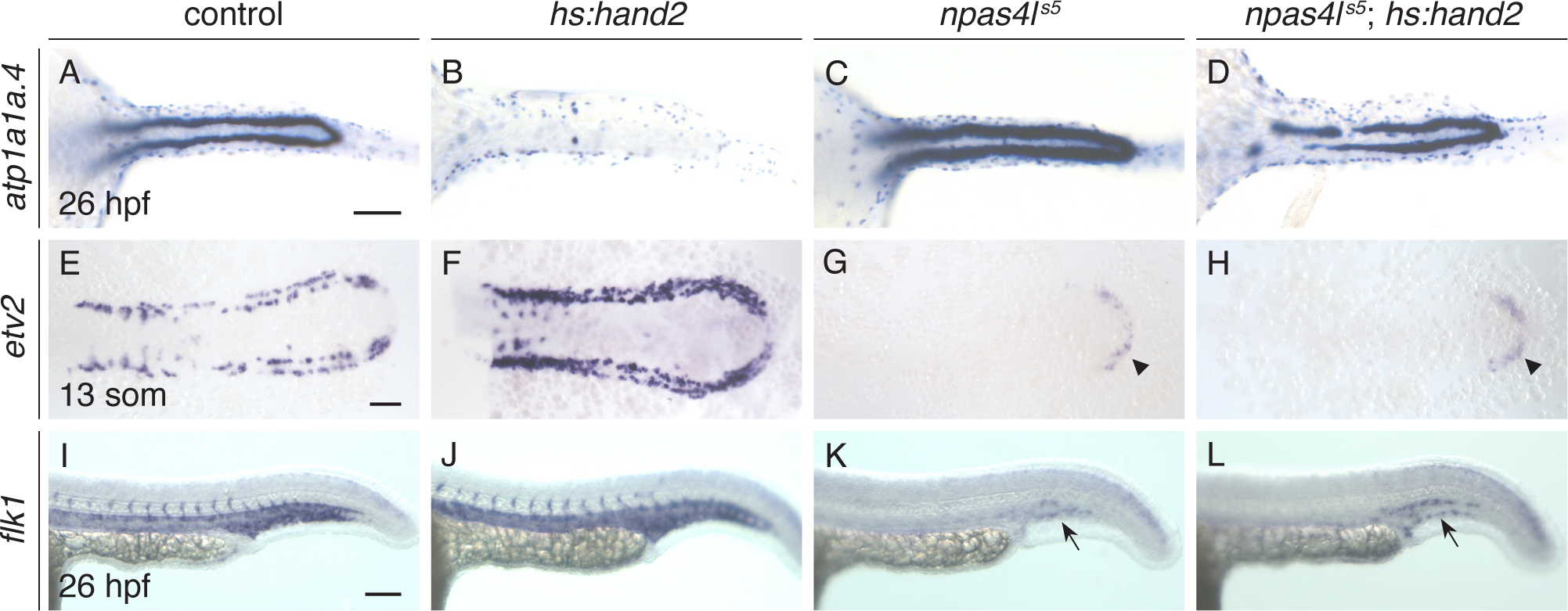
*hand2* overexpression requires *npas4l* function to inhibit pronephron formation. *In situ* hybridization demonstrates *atp1a1a.4* (A-D) expression throughout the pronephron tubules, *etv2* (E-H) expression in vessel progenitors, and *flk1* (I-L) expression in the vasculature. Dorsal views, anterior to the left, of wild-type (B,F,J) and *npas4l^s5^* mutant embryos (D,H,L) carrying *hs:hand2* were compared with their corresponding wild-type control (A,E,I) and *npas4l^s5^* mutant (C,G,K) nontransgenic siblings. Heat shock was performed at tailbud in all embryos shown. Compared to wild-type control embryos (A), *atp1a1a.4* expression in the pronephron tubules was expanded in *npas4l^s5^* mutants (C; n=17/19). Unlike the dramatically decreased *atp1a1a.4* expression in *hs:hand2* embryos (B), *atp1a1a.4* expression in the pronephron of *npas4l^s5^*;*hs:hand2* embryos is only relatively mildly reduced (D; n=34/38). Compared to wild-type (E), *etv2* expression appears more intense and is observed in an increased number of cells, especially within the territory normally occupied by the IM, in *hs:hand2* embryos (F). *etv2* (G,H) and *flk1* (K,L) expression are largely absent in both *npas4l^s5^*mutants (G,K), and *npas4l^s5^*;*hs:hand2* embryos (H,L). Note the faint *etv2* expression in the tailbud of the *npas4l^s5^* mutants (arrowheads) and *flk1* expression in the posterior trunk of the *npas4l^s5^* mutants (arrows) are consistent with prior reported phenotypes of *npas4l* mutants (Liao et al., 1997; Reischauer et al., 2016; Sumanas et al., 2005). n=65 (A), 129 (B), 19 (C), 38 (D), 119 (E), 118 (F), 37 (G), 33 (H), 19 (I), 8 (K), 6 (L). Scale bars: 100 μm.

### *npas4l* can inhibit pronephron formation independent of *hand2* function

To further examine whether *npas4l* may function downstream of *hand2* in pronephron development, we next sought to determine the capacity of *npas4l* activity in the absence of *hand2* function. We found that overexpression of *npas4l* was sufficient to inhibit pronephron tubule formation (Fig. 3A,C). Importantly, *npas4l* retained this capacity in the absence of *hand2* function (Fig. 3B,D), consistent with *npas4l* inhibiting pronephron formation downstream of or in parallel with *hand2* function.

**Figure 3.**
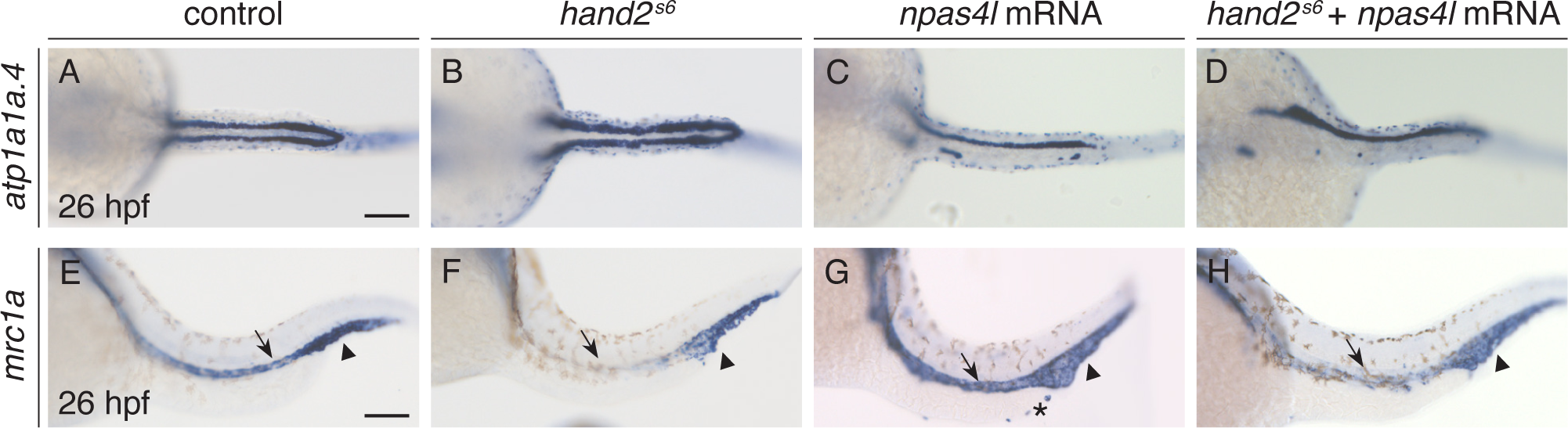
*npas4l* overexpression inhibits pronephron formation in the absence of *hand2* function. Dorsal views, anterior to the left, of wild-type control embryos (A,E), *hand2^s6^* mutant embryos (B,F), and wild-type control (C,G) and *hand2^s6^* mutant embryos (D,H) injected with *npas4l* mRNA, all at 26 hpf. *In situ* hybridization demonstrates *atp1a1a.4* (A-D) expression throughout the pronephron tubules, and *mrc1a* (E-H) expression in the posterior cardinal vein (arrow) and posterior blood island (arrowhead) (Stachura & Traver, 2016). Compared to wild-type (A), pronephron gene expression is expanded in *hand2^s6^*mutants (B) and greatly reduced in *npas4l*-overexpressing embryos (C,D). The extent of the defects seen in *npas4l*-overexpressing embryos was comparable between injected *hand2^s6^* mutants (D; n=10/28 with reduced pronephron as shown) and injected wild-type control embryos (C; n=26/79 with reduced pronephron as shown). Compared to wild-type (E), *mrc1a* gene expression in the posterior cardinal vein (arrow) is greatly reduced in both uninjected and *npas4l*-overexpressing *hand2^s6^* mutants (F,H) but is increased in intensity in *npas4l*-overexpressing embryos (G). *npas4l* mRNA overexpression also resulted in ectopic *mrc1a* expression along the yolk extension (asterisk in G; n=41/75 with ectopic expression). Although higher doses of *npas4l* mRNA led to stronger phenotypes than shown here, they also increased embryonic lethality or severe deformation. Therefore, the dose of *npas4l* mRNA injected was limited to 5-14.5 ng so that overall development was not severely impacted which likely limited the penetrance and expressivity of phenotypes displayed. n=28 (A), 11 (B), 79 (C), 28 (D), 38 (E), 13 (F), 75 (G), 19 (H). Scale bars: 100 μm.

In addition to the role of *hand2* in inhibiting IM and pronephron formation, we previously found that *hand2* mutants fail to express *mrc1a* in the cardinal vein (Fig. 3E,F), which was thought to be caused by the lack of LVPs (Kohli et al., 2013; Perens et al., 2016). Because *hand2* mutants lack *npas4l* expression in LVPs (Fig. 1H), we wanted to determine whether *npas4l* overexpression could rescue the cardinal vein defect in *hand2* mutants. While *npas4l* overexpression could elevate the expression of *mrc1a* in wild-type embryos (Fig. 3G), it could not rescue the *mrc1a* expression defect in the posterior cardinal vein of *hand2* mutants (Fig. 3H). Thus, our data further suggest that *hand2* is required for certain elements of venous differentiation, such as the expression of vein-specific markers.

In sum, while *npas4l* cannot promote vein formation in the absence of *hand2*, it can inhibit pronephron formation. Nevertheless, it remains unclear where and when *npas4l* inhibits pronephron formation. Up to this point, our data suggest that *npas4l* acts downstream of *hand2* and *osr1* to promote LVP differentiation and prevent excess pronephron formation from cells at the interface between the IM and *hand2*/*osr1*-expressing territories. However, because *npas4l* is also expressed along the medial border of the IM, the possibility remains that *npas4l* has an additional, independent function separate from *hand2* or *osr1* in suppressing pronephron formation.

### Removal of *npas4l* function restores pronephron development in *osr1* mutants, but loss of *osr1* fails to rescue *npas4l* mutant vessel defects

A remarkable feature of the role of *hand2* in inhibiting pronephron development is that removal of *hand2* function can rescue the pronephron defects caused by *osr1* loss-of-function (Perens et al., 2021; Perens et al., 2016). Specifically, in *osr1* mutants, expression of glomerular precursor markers is absent and the pronephron tubules are thin and lack *cdh17* expression at the proximal end of the tubule (Fig. 4A,B); each of these phenotypes is partially suppressed by *hand2* loss-of-function (Drummond et al., 2022; Perens et al., 2021). Conversely, removal of *osr1* function can partially rescue the LVP deficits caused by *hand2* loss of function (Perens et al., 2021). Because *hand2* is expressed with *osr1* in the lateral territory flanking the IM (Fig. 1A,B; (Perens et al., 2016)), these findings support a model in which *hand2* and *osr1* function together to balance the contribution of progenitors at the lateral IM border to pronephron and LVP fates. To help further understand the function of *npas4l* and compare the roles of *npas4l* and *hand2* in pronephron fate inhibition, we sought to determine if *npas4l* and *hand2* exhibit similar genetic interactions with *osr1*.

**Figure 4.**
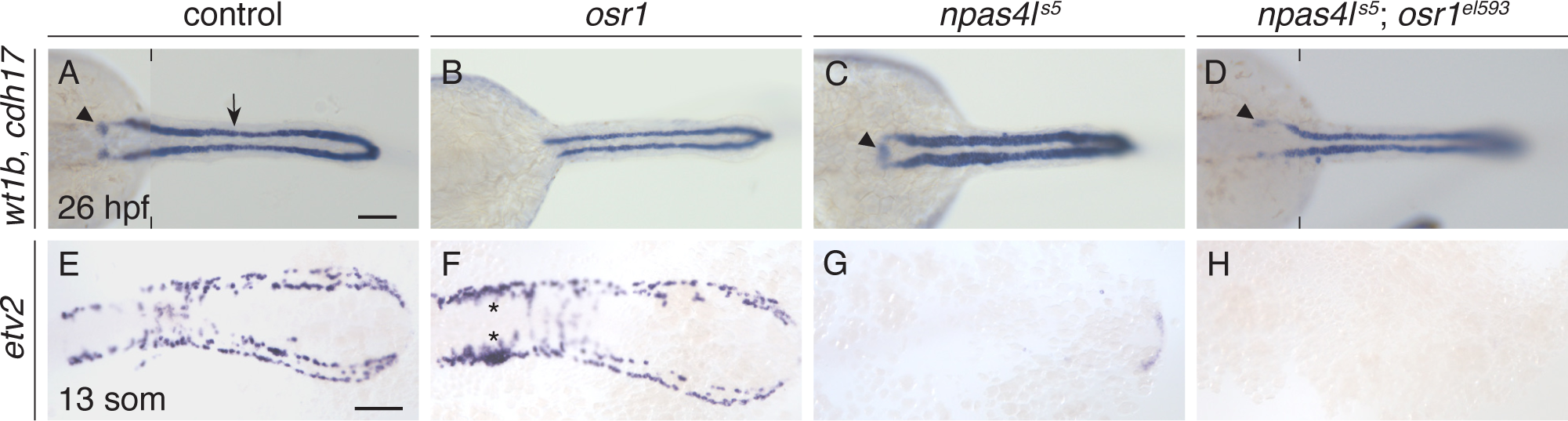
Pronephron defects in *osr1^el593^* mutants are partially suppressed by loss of *npas4l* function. Dorsal views, anterior to the left, of wild-type control (A,E), *osr1^el593^* mutant (B,F), *npas4l^s5^* mutant (C,G) and *npas4l^s5^*;*osr1^el593^* double mutant (D,H) embryos at 26 hpf (A-D) and 13 som (E-H). *In situ* hybridization shows expression of *wt1b* in glomerular precursors (arrowhead), *cdh17* throughout the pronephron tubules (arrow) (A-D), and *etv2* in vessel progenitors (E-H). Compared with wild-type controls (A), pronephron glomerular expression is absent and tubule expression is thin and shortened anteriorly in *osr1^el593^* mutants (B), expanded in *npas4l^s5^* mutants (C); and relatively similar to wild-type in *npas4l^s5^*;*osr1^el593^* double mutants (D; n=8; 75% with normal tubules and *wt1b* glomeruli expression; 25% with normal tubules and absent *wt1b* glomeruli expression). In contrast, compared with wild-type controls (E), *etv2* vessel progenitor expression is unaffected in *osr1* mutants (F; expression was expanded in the midbody region (asterisks), as previously reported (Perens et al., 2021)), but is mostly absent in both *npas4l^s5^* mutants (G) and *npas4l^s5^*;*osr1^el593^*double mutants (H). n=112 (A), 49 (B), 43 (C), 8 (D), 64 (E), 19 (F), 21 (G), 6 (H). Scale bars: 100 μm.

First, we found that *osr1* loss-of-function could suppress the excess pronephron formation seen in *npas4l* mutants. In *npas4l* mutants, we observed expanded glomerular precursors and pronephron tubules (Fig. 4C), similar to defects seen in *hand2* mutants (Perens et al., 2016) and consistent with previously characterized *npas4l* mutant pronephron phenotypes (Majumdar & Drummond, 1999; Mattonet et al., 2022). Remarkably, in *npas4l*;*osr1* double mutants, glomerular precursors and pronephron tubules appeared most similar to wild-type embryos (Fig. 4A-D). Thus, like the interaction between *hand2* and *osr1*, the increased pronephron formation caused by *npas4l* loss-of-function could compensate for the decreased pronephron formation due to *osr1* loss-of-function. Is the same interaction observed for vessel progenitor formation? Notably, in contrast to the interaction observed between *hand2* and *osr1* (Perens et al., 2021), none of the extensive vessel progenitor defects observed in *npas4l* mutants, including the absence of LVPs, were restored by removing *osr1* function (Fig. 4E-H). Thus, *npas4l* and *hand2* do not exhibit identical genetic interactions with *osr1*.

In sum, in *hand2*;*osr1* double mutants, it appears as though when the excess pronephron caused by *hand2* loss-of-function is reduced to near wild-type levels by *osr1* loss-of-function, there is a corresponding increase/rescue of the LVPs lacking in the *hand2* single mutant (Perens et al., 2021). In contrast, in *npas4l*;*osr1* double mutants, when the excess pronephron caused by *npas4l* loss-of-function is reduced by *osr1* loss-of-function (Fig. 4A-D), there is not a concomitant increase in vessel progenitor formation (Fig. 4E-H). Combined with our prior observation that, unlike *hand2* and *osr1*, *npas4l* is expressed along both the lateral and medial borders of the IM (Fig. 1A,B), these observations raise the possibility that some of the excess pronephron observed in *npas4l* mutants may be derived from a different source than the excess pronephron observed in *hand2* mutants.

### *npas4l* mutants exhibit increased IM outside of the *hand2*-expressing territory

Mattonet and colleagues demonstrated that the number of pronephron tubule cells is increased in *npas4l* mutants at 24 hours post fertilization (hpf) (Mattonet et al., 2022). Morphogenesis of the pronephron tubules from the IM initiates at 14 som (approximately 16 hpf) with lumen formation occurring between 20-22 som (approximately 19-20 hpf) (Gerlach & Wingert, 2014), and Mattonet and colleagues did not observe a qualitative change in the population of pronephron progenitors (as labeled by Pax2a) in *npas4l* mutants at 12 hpf (Mattonet et al., 2022). Thus, the timing and origin of the excess pronephron observed in *npas4l* mutants remained unclear.

Previously, we quantified the change in Pax2a^+^ cells in *hand2* and *osr1* mutants at 10 som (approximately 14 hpf) (Perens et al., 2021; Perens et al., 2016). To more rigorously interrogate the *npas4l* IM phenotype, here we performed similar analysis of *npas4l* mutants. We found a significant increase in the number of Pax2a^+^ cells within the IM of *npas4l* mutants at 11 som (Fig. 5A,B,D). Thus, the excess pronephron in *npas4l* mutants originates prior to pronephron morphogenesis, and we conclude that, like *hand2*, *npas4l* inhibits excess pronephron development by limiting IM formation.

**Figure 5.**
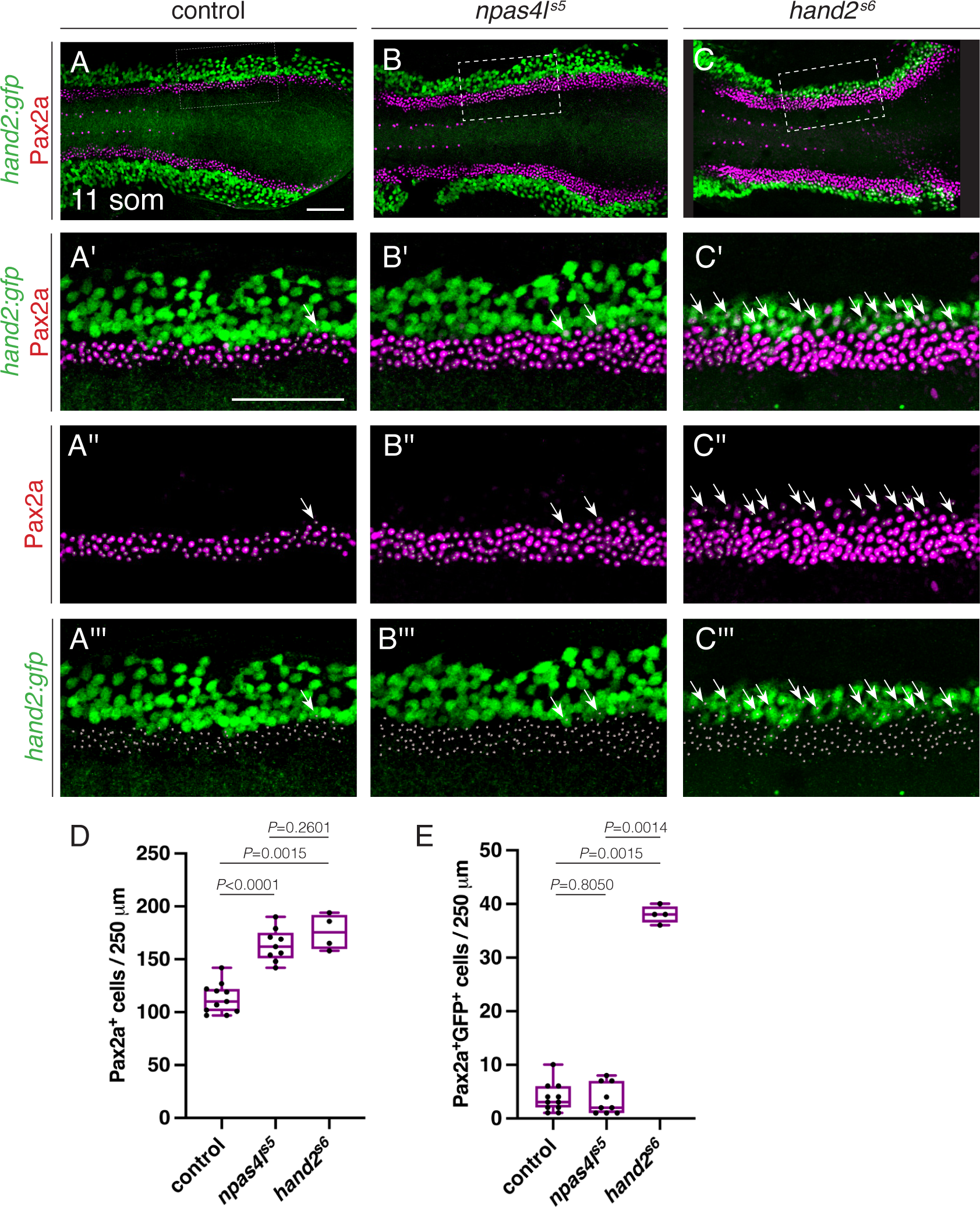
*npas4l* inhibits IM formation. Immunofluorescence for Pax2a and GFP in wild-type (A), *npas4l^s5^*mutant (B), and *hand2^s6^* mutant (C) embryos carrying *Tg*(*hand2:EGFP*); dorsal views, anterior to the left, of three-dimensional reconstructions at 11 som. (A’-C’’’) Magnification of regions from (A-C) used for quantification of the numbers of Pax2a^+^ cells. 250 μm stretches of the IM were analyzed per embryo. The length of the analyzed region varies within the boxed region due to mild curvature of the embryo in the z-axis. White dots indicate Pax2a^+^ nuclei and arrows indicate examples of Pax2a^+^GFP^+^ cells. Scale bars: 100 µm. (D,E) Quantification of the numbers of Pax2a^+^ cells (D) and Pax2a^+^GFP^+^ cells (E) per 250 μm of IM in wild-type control, *npas4l^s5^*, and *hand2^s6^* embryos demonstrates an increase in IM cells in *npas4l^s5^*and *hand2^s6^* embryos, but an increase in Pax2a^+^GFP^+^ cells only in *hand2^s6^* embryos. Boxes represent interquartile range, central line marks the median, and whiskers indicate maximum and minimum values. *P* values were calculated using non-parametric Mann–Whitney *U*-tests. n=11 (A), 9 (B), 4 (C).

Furthermore, previously we showed that *hand2* constrains the extent of pronephron formation by inhibiting IM formation within the laterally adjacent *hand2*-expressing cells (Perens et al., 2016). Unlike in *hand2* mutants (Fig. 5C,E), however, excess Pax2a^+^ cells in *npas4l* mutants were observed outside of the *hand2*-expressing territory (Fig. 5B,E). That is, while *npas4l* and *hand2* mutants have comparable amounts of Pax2a^+^ cells in the posterior mesoderm (Fig. 5D), only *hand2* mutants have a significantly increased number of Pax2a^+^ cells within the lateral *hand2*-expressing territory (Fig. 5E). Thus, *npas4l* inhibits excess IM formation in a portion of the posterior mesoderm distinct from the *hand2*-expressing territory.

Consistent with non-overlapping roles for *npas4l* and *hand2* in inhibiting excess IM formation, *hand2*;*npas4l* double mutants have an increased amount of IM beyond that seen in either single mutant. More specifically, in the double mutants, we observed a qualitative increase in the width of the IM based on *pax2a* (Fig. 6A-D) and *lhx1a* (Fig. S4) expression, and a quantitative increase in Pax2a^+^ cells (Fig. 6E-I). Additionally, in analyzing these embryos, we also noted that *hand2* mutants and *hand2*;*npas4l* double mutants exhibited a particular IM phenotype distinct from that observed in *npas4l* mutants. In *hand2* mutants and *hand2*;*npas4l* double mutants, we frequently observed less intense expression of IM markers near the lateral edge of the IM (arrows in Fig. 6C’,D’,G’,H’, Fig. S4C’,D’). This variation in intensity was apparent based on both *pax2a* and *lhx1a* expression (Fig. 6A-D, Fig. S4) and Pax2a localization (Fig. 6E-H). Presumably, this represents the ectopic IM found in the *hand2*-expressing territory in *hand2* mutants (Fig. 5C) but not in *npas4l* mutants (Fig. 5B). This difference again highlights the apparently divergent origins of increased IM in the *hand2* and *npas4l* mutants. Together, these findings suggest that in addition to a role for *npas4l* inhibiting IM specification downstream of *hand2* and *osr1* at the lateral IM border, *npas4l* is also required for inhibition of excess IM specification independent from *hand2* at the medial border of the IM.

**Figure 6.**
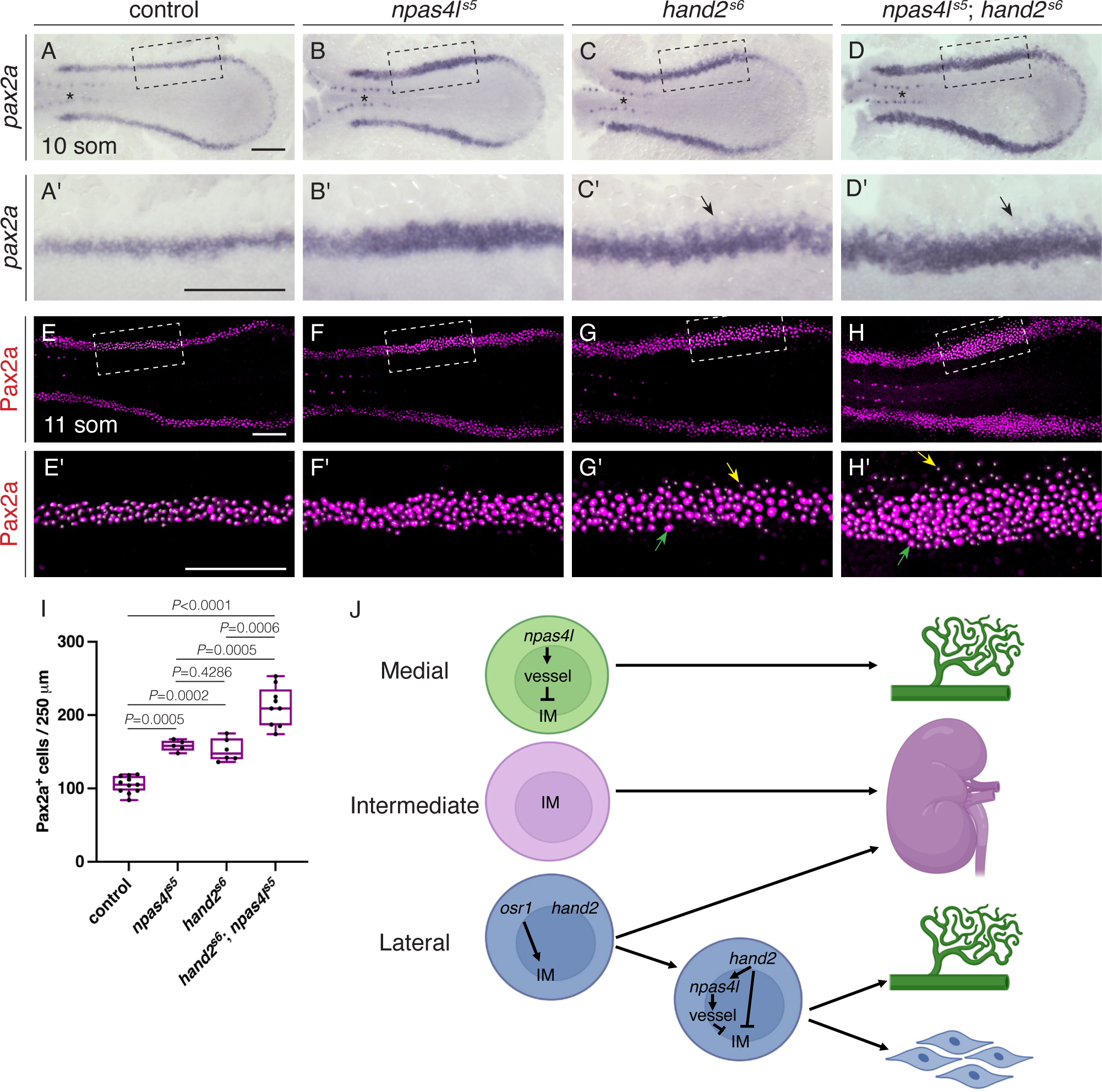
IM defects in *npas4l^s5^*mutants are enhanced by loss of *hand2* function. (A-H) Dorsal views, anterior to the left, of the posterior mesoderm. (A-D) *In situ* hybridization shows expression of *pax2a* in the IM (arrows) at 10 som. Compared with wild-type (A), expression is widened in *npas4l^s5^* mutants (B) and *hand2^s6^* mutants (C) and widened further in *hand2^s6^*;*npas4l^s5^* double mutants (D) Expression in the overlying spinal neurons (asterisk) is unaffected. Unlike the width of the IM, we did not observe a noticeable change in the proximal-distal length of the IM in mutant embryos. (A’-D’) Magnification of boxed regions in (A-D). Faint lateral staining (arrows) was frequently observed in *hand2^s6^* mutant (C’) and *hand2^s6^*;*npas4l^s5^* double mutant (D’) embryos. n=60 (A), 17 (B), 21 (C), 7 (D). (E-H) Three-dimensional reconstructions of Pax2a immunofluorescence in the IM of wild-type (E), *npas4l^s5^* mutant (F), *hand2^s6^* mutant (G) and *hand2^s6^*;*npas4l^s5^* double mutant (H) embryos at 11 som. (E’-H’) Magnification of boxed 250 µm long regions used for counting Pax2a^+^ cells. White dots indicate Pax2a^+^ nuclei. In *hand2^s6^* mutant and *hand2^s6^*;*npas4l^s5^* double mutant embryos, intensity of Pax2a^+^ staining varied from strong (for example, green arrows) to weak (for example, yellow arrows). As seen with *in situ* hybridization, the fainter Pax2a^+^ levels were typically observed at the lateral edge in *hand2^s6^* mutants (G’) and *hand2^s6^*;*npas4l^s5^*double mutants (H’). n=11 (E), 5 (F), 6 (G), 9 (H). Scale bars: 100 µm. (I) Quantification of the numbers of Pax2a^+^ cells per 250 µm of IM in the indicated genotypes demonstrates an increase in IM cells in *hand2^s6^*;*npas4l^s5^*double mutant embryos exceeding that seen in either single mutant. Boxes represent interquartile range, central line marks the median, and whiskers indicate maximum and minimum values. *P* values were calculated using non-parametric Mann–Whitney *U-*tests. (J) Model of organ field formation in the lateral posterior mesoderm in zebrafish. In the most medial territory, *npas4l* promotes expression of vessel progenitor genes (labeled “vessel”; including *etv2*, *lmo2*, and *tal1*), which inhibit IM. The intermediate territory expresses IM-associated genes (including *lhx1a* and *pax2a*). Initially, the most lateral territory expresses both *osr1* and *hand2* (Perens et al., 2016), and *osr1* is required for forming the full complement of IM (Perens et al., 2021). Later, as *osr1* expression recedes (as shown in (Perens et al., 2021)), *hand2* expression promotes *npas41* expression, which inhibits further IM gene expression and results in the generation of the LVPs. The remainder of the lateral territory continues to express *hand2* and gives rise to other lateral plate mesoderm derivatives, including the mesothelium (Prummel et al., 2022).

## DISCUSSION

Our data provide new insights into the strategies implemented to divide the posterior mesoderm into defined organ progenitor territories. In particular, our findings highlight how the dimensions of the IM are defined by factors that promote neighboring vessel progenitor fates. We find that *npas4l* is required to inhibit excess IM formation. Based on our analysis, we propose that *npas4l* is required to repress IM fate at both the lateral and medial IM boundaries, and, at the lateral boundary, *hand2* and *osr1* act genetically upstream of *npas4l* to regulate a decision between IM and vessel progenitor fates. Given that Mattonet and colleagues observed ectopic Pax2a expression in *npas4l*-expressing cells lacking *npas4l* function (Mattonet et al., 2022), the increased IM generated in *npas4l* mutants appears to result from a cell fate conversion in which mesodermal progenitors normally fated to give rise to vessel progenitors instead become pronephron progenitors within the IM. Altogether, these findings suggest a tight coupling of IM and vessel progenitor fates in the posterior mesoderm: transcription factors that promote vessel fate (*hand2*, *npas4l*) and those that promote IM fate (*osr1*) each inhibit the alternative fate. This mechanism of mutual repression ensures precise gene expression boundaries between these neighboring organ progenitor territories. Interestingly, similar reciprocal repression has been observed between lineage-defining genes during the vascular versus myogenic cell fate decision in the dermomyotome (Lagha et al., 2009).

Combined with our prior analyses of *hand2* and *osr1* (Perens et al., 2021; Perens et al., 2016), our findings suggest different gene regulatory networks are employed to coordinate IM and vessel progenitor fates in distinct territories along the medial-lateral axis of the zebrafish posterior mesoderm (Fig. 6J). Based on our gene expression analysis (Fig. 1A), each side of the lateral posterior mesoderm is initially divided into three territories along the medial-lateral axis. An intermediate territory, marked by *pax2a* and *lhx1a* expression, presumably gives rise to the pronephon. However, as we previously proposed (Perens et al., 2021), *osr1* function is required to generate the full complement of IM and, subsequently, the complete pronephron. Meanwhile, the medial territory is demarcated by *npas41* expression. Our data here suggest that *npas4l* both inhibits IM and promotes vessel progenitor fate decision in this location. Last, *hand2* and *osr1* expression mark the lateral territory. Some progenitors within this lateral territory continue to express *hand2* and migrate medially (Yin et al., 2010), giving rise to derivatives that include the mesothelium (Prummel et al., 2022). Notably, *hand2* is an essential and potent inhibitor of IM fate within this territory; we have only observed ectopic Pax2a expression in this territory in *hand2* mutants, but not in *npas4l* loss-of-function or *osr1* overexpression scenarios.

Importantly, a fourth territory comprised of vessel progenitors (the LVPs) expressing *etv2*, *tal1*, and *npas4l* later arises at the boundary between the intermediate and lateral territories (Fig. 1B, Fig. S1; (Kohli et al., 2013; Perens et al., 2016)). The origin of these LVPs remains unclear. Previously, we proposed that the lateral territory initially gives rise to additional IM and later to LVPs (Perens et al., 2021). Our data here suggest that in this territory, *hand2* and *osr1* influence these cell fate decisions through regulation of *npas4l* (Fig. 6J). In this model, progenitors within this territory are initially promoted to the IM fate under the control of *osr1*, which also inhibits *npas4l* expression. Later, as *osr1* expression decreases (Perens et al., 2021), *hand2* via *npas4l* inhibits further IM formation and instead instructs the progenitors to give rise to the LVPs.

Our studies highlight the extensive connections between the specification of kidney and vessel lineages. Perhaps most intriguing is the possibility of a common bipotential progenitor that gives rise to both the kidney and vessel lineages. Within the context of our model for the patterning of the zebrafish posterior mesoderm, the origin of the LVPs and *osr1*-dependent IM remain unclear. We propose that each could arise from the lateral *hand2-* and *osr1*-expressing territory (Fig. 6J). Alternatively, one or both could arise from the *pax2a*-expressing intermediate territory. Interestingly, recent single cell profiling of endothelial progenitors in zebrafish identified a small population that expresses IM markers such as *pax2a* and *osr1* (Mattonet et al., 2022; Sahai-Hernandez et al., 2023), further raising the possibility of a shared progenitor. Additionally, although not direct evidence of a bipotential progenitor, it is intriguing that genetic fate mapping of *pax2a*-expressing descendants in zebrafish and *Osr1*-expressing descendants in mouse labeled both kidney and vasculature (Mattonet et al., 2022; Mugford et al., 2008). Ultimately, high-resolution fate mapping will be necessary to determine the existence of a bipotential progenitor, as well as to determine the fates of the different posterior mesoderm territories and how they are altered in different genetic backgrounds. Whether there is a bipotential progenitor or not, mounting evidence continues to suggest a strong interconnection between these fate choices. For example, as mentioned above, Mattonet and colleagues found that *npas4l*-expressing cells express *pax2a* and contribute to the pronephron when lacking *npas4l* function (Mattonet et al., 2022). Furthermore, we have observed that *hand2* overexpression only elevates vessel progenitor gene expression in the territory that normally forms the IM (Fig. 2F; (Perens et al., 2016)), suggesting that the IM appears especially susceptible to expressing markers of vessel progenitor fate.

How the Hand2, Osr1, and Npas4l transcription factors regulate IM and vessel progenitor gene expression also remains unknown. It is possible that each of these transcription factors independently regulates the transcription of IM factors, such as *pax2a* and *lhx1a*, and vessel progenitor factors, such as *etv2*. It is also possible that Hand2 and Osr1 only indirectly impact IM and/or vessel gene expression. Here we find that the ability of *hand2* to inhibit pronephron formation was drastically diminished by *npas4l* loss-of-function, suggesting that Hand2 does not directly impact IM gene expression, but rather does so via factors that promote vessel progenitor fate, including Npas4l. Interestingly, by comparing transcriptomes and chromatin accessibility between wild-type and *hand2* crispant LPM cells, Komatsu and colleagues recently suggested that Hand2 represses IM and pronephron gene expression within the LPM by repressing chromatin accessibility of their associated regulatory elements (Komatsu et al., 2023). Whether this effect is direct, however, is undetermined. Additionally, it is unclear if each transcription factor acts as a transcriptional activator and/or repressor when regulating IM and vessel progenitor fates. In the case of Osr1, in *Xenopus*, Osr1 fused to the Engrailed repressor domain, but not the E1A activation domain, was sufficient to induce ectopic IM gene expression (Tena et al., 2007). Nevertheless, additional studies will be necessary to determine the molecular mechanisms each transcription factor implements to impact gene expression involved in these fate decisions.

Understanding the genetic regulatory networks that coordinate the development of kidneys and vessels has the potential to provide valuable insights into the etiologies of congenital anomalies of the kidney and urinary tract (CAKUT) and could offer valuable guidance for enhancing the effectiveness of kidney stem cell and regenerative technologies. The relatively high degree of familial aggregation of CAKUT suggests a significant genetic component, and disease-causing mutations have already been identified in dozens of genes, including many genes required for IM development, such as *PAX2, OSR1*, and *HNF1β* (Fillion et al., 2017; Kolvenbach et al., 2023; Sanna-Cherchi et al., 2018; Thomas et al., 2011). Unraveling the gene regulatory networks that help define the IM, could expand the spectrum of candidate genes associated with CAKUT. Regarding stem cell and regenerative techniques, it is particularly intriguing to note that kidney organoids generated from putative IM possess endothelial cells (Takasato et al., 2015). Some have postulated that this outcome is due to an off-target effect during the directed differentiation process (Ryan & Cleaver, 2022). Whether this outcome is an off-target effect or is due to an inherent potential for IM to give rise to endothelial cells, our studies examining the genetic and lineage relationships between kidney and endothelial progenitors should yield valuable insights necessary for understanding and refining protocols implemented during the in vitro generation of kidney lineages.

## Acknowledgements

We thank members of the Yelon lab for valuable discussions, A. Yarbrough and the UCSD Animal Care Program for zebrafish care. Fig. 6J was created with BioRender.com.

## Data availability

Data will be made available on request.

## Author contributions

Conceptualization: E.A.P., D.Y.; Data curation: E.A.P., D.Y.; Formal analysis: E.A.P., D.Y.; Funding acquisition: E.A.P., D.Y.; Investigation: E.A.P.; Methodology: E.A.P., D.Y.; Project administration: D.Y; Resources: E.A.P., D.Y.; Supervision: D.Y.; Validation: E.A.P., D.Y.; Visualization: E.A.P., D.Y.; Writing - original draft: E.A.P.; Writing - review and editing: E.A.P., D.Y;.

---

**Supplementary Figure S1.**
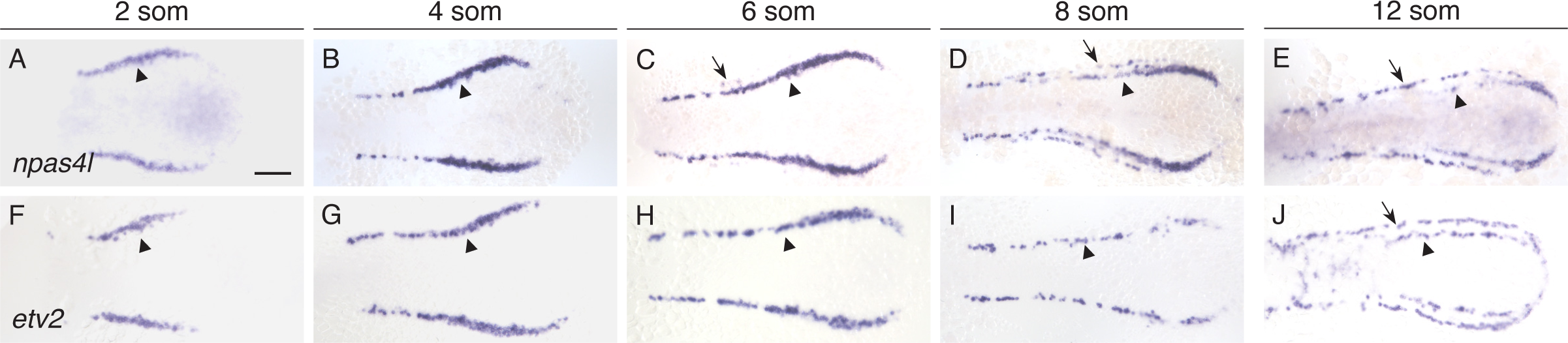
*npas4l* expression dynamics precede *etv2* in the posterior mesoderm. Dorsal views, anterior to the left, of *in situ* hybridization in wild-type embryos at 2 som (A,F), 4 som (B,G), 6 som (C,H), 8 som (D,I), and 12 som (E,J). (A-E) *npas4l* is transiently expressed in medial vessel progenitors (arrowheads). Starting at 6 som, expression begins to recede in an anterior to posterior manner, until at 12 som only a few *npas4l*-expressing medial progenitors persist at the posterior. *npas4l* expression in LVPs (arrows) is observed in a small number of cells in an anterior portion of the posterior mesoderm as early as 6 som (C;, n=156; 19% with LVP expression), and expression increases in an anterior to posterior manner such that embryos at 8 som (D; n=63; 95%) and 12 som (E; n=32; 100%) typically express *npas4l* throughout the LVPs. (F-J) In contrast, *etv2* remains expressed in relatively medial vessel progenitors (arrowheads) between 2 and 12 som and is expressed in LVPs (arrows) starting at 11 som (Perens et al., 2016). n=65 (A), 79 (B), 156 (C), 63 (D), 32 (E), 8 (F), 11 (G), 9 (H), 11 (I), 19 (J). Scale bar: 100 μm.

**Supplementary Figure S2.**
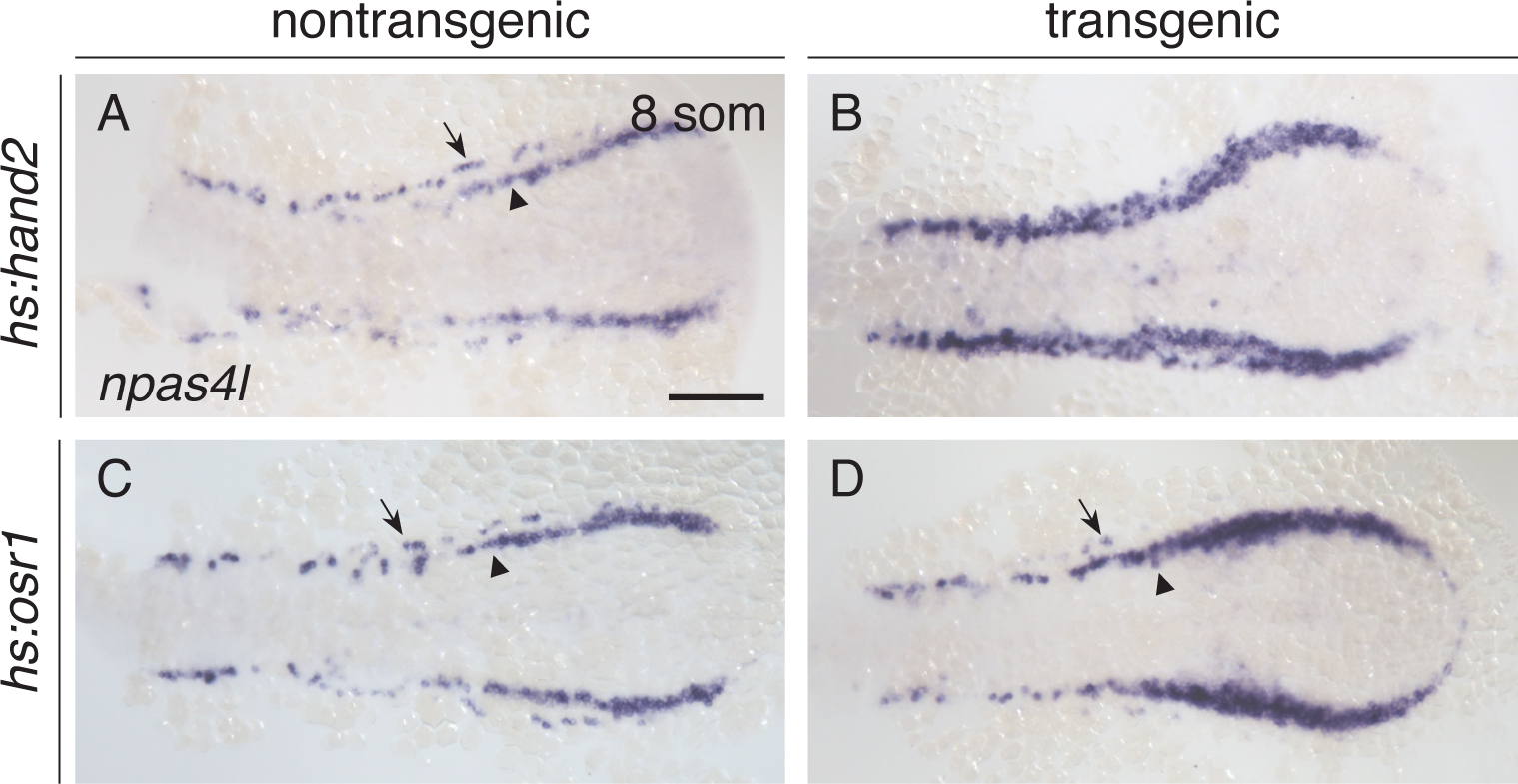
***hand2* and *osr1* overexpression affect the development of *npas4l*-expressing vessel progenitors in the posterior mesoderm.** *In situ* hybridization depicts *npas4l* expression in nontransgenic controls (A,C), *Tg*(*hsp70:hand2-2A-mcherry*) (*hs:hand2*) (B), and *Tg*(*hsp70:osr1-t2A-BFP*) (*hs:osr1*) (D) embryos; dorsal views, anterior to the left, at 8 som. (A,C) *npas4l* is expressed in medial (arrowheads) and lateral (arrows) territories on each side of the nontransgenic control embryos heat shocked at tailbud stage. When *hand2* is overexpressed in *hs:hand2* embryos (B), the number of cells expressing *npas4l* is increased, especially within the territory normally occupied by the IM. It is unclear whether this expression represents an increase in the medial or lateral vessel progenitor populations, since no currently available markers can distinguish these two groups of progenitors. When *osr1* is overexpressed in *hs:osr1* embryos (D) display decreased *npas4l* expression in lateral territories (n=56). Most embryos (97%) displayed no LVP expression, while a minority (3%) displayed a small amount of *npas4l* expression at the anterior aspect of the lateral territories (D, arrow). In contrast, expression of *npas4l* in medial territories (arrowhead) was expanded. These phenotypes are comparable to those observed with *etv2* expression in embryos overexpressing *osr1*, and we previously demonstrated that the increased *etv2* expression was medial to *pax2a* expression, consistent with inhibition of LVP formation (Perens et al., 2021). n=78 (A), 88 (B), 29(C), 56 (D). Scale bar: 100 μm.

**Supplementary Figure S3.**
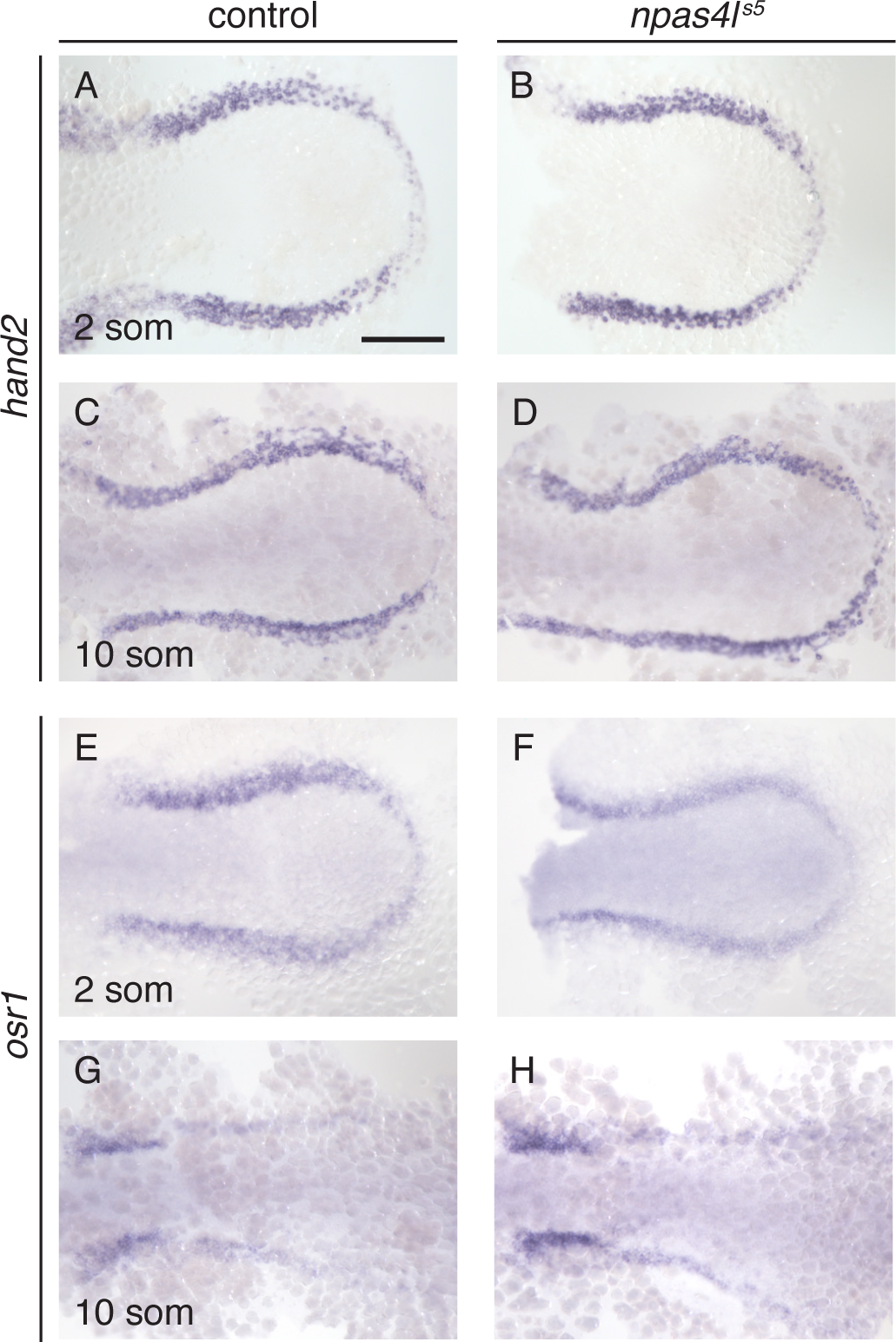
Expression patterns of *hand2* and *osr1* appear unaffected by loss of *npas4l* function. (A-H) *In situ* hybridization depicts expression of *hand2* (A-D) and *osr1* (E-H) in the posterior mesoderm of wild-type (A,C,E,G) and *npas4l^s5^* mutant (B,D,F,H) embryos; dorsal views, anterior to the top, of the posterior portion of the embryo at 2 som (A,B EF) and 10 som (C, D,G,H). Loss of *npas4l* function does not appear to affect *hand2* (A-D) or *osr1* (E-H) expression. n=70 (A), 24 (B), 18 (C), 5 (D), 17 (E), 7 (F), 105 (G), 37 (H). Scale bar: 100 μm.

**Supplementary Figure S4.**
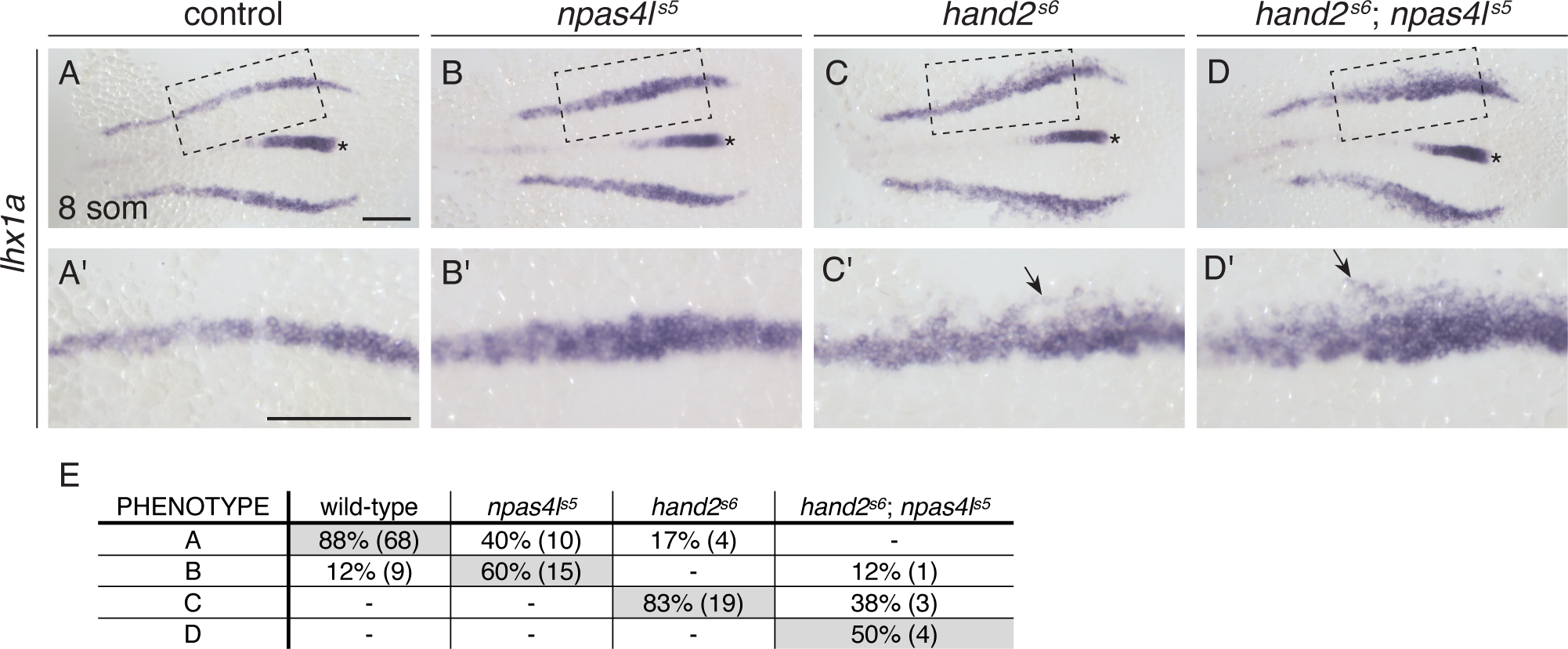
IM defects in *npas4l^s5^* mutants are enhanced by *hand2* loss of function. (A-D) Dorsal views, anterior to the left, of the posterior mesoderm at 10 som. *In situ* hybridization shows expression of *lhx1a* in the IM. Compared with wild-type (A), expression is widened in *npas4l^s5^*mutants (B) and *hand2^s6^* mutants (C) and is widened further in *npas4l^s5^*;*hand2^s6^* (D) embryos. Expression in the notochord (asterisk) is unaffected. (A’-D’) Magnification of boxed regions in (A-D). As with *pax2a* expression and Pax2a localization (Fig. 6), fainter expression was observed at the lateral edge of the *hand2^s6^*mutant (C’) and *hand2^s6^*;*npas4l^s5^* double mutant (D’) embryos (arrows). n=77 (A), 25 (B), 23 (C), 8 (D). Scale bars: 100 µm. (E) Table comparing phenotypes of *lhx1a* expression in the IM represented in A-D to the corresponding genotypes. Values represent the percentage of embryos with the corresponding genotype that display a particular phenotype; n values are in parentheses. The phenotypes included (A) narrow (approximately 2-3 cells wide) wild-type-like *lhx1a* expression in the IM, (B) widened (approximately 4-5 cells wide) *lhx1a* expression without faint lateral expression, (C) widened (approximately 4-5 cells wide) *lhx1a* expression with faint lateral expression, and (D) dramatically widened (approximately 6-9 cells wide) *lhx1a* expression with faint lateral expression. Shaded boxes indicate case in which the phenotype is the one depicted for the corresponding genotype in A-D.

